# Overcoming ploidy barriers: the role of triploid bridges in the genetic introgression of *Cardamine amara*

**DOI:** 10.1101/2024.10.08.617200

**Authors:** P Bartolić, A Voltrová, L Macková, G Šrámková, M Šlenker, T Mandáková, N Padilla García, K Marhold, F Kolář

**Affiliations:** Department of Botany, Faculty of Science, Charles University in Prague, Benátská 2, CZ-128 01 Prague, Czechia; Institute of Botany, Plant Science and Biodiversity Centre, Slovak Academy of Sciences, SK-845 23 Bratislava, Slovakia; Department of Experimental Biology, Faculty of Science, Masaryk University, Brno, Czechia; Central European Institute of Technology, Masaryk University, Brno, Czechia; Departamento de Botánica y Fisiología Vegetal, University of Salamanca, 37007 Salamanca, Spain; Institute of Botany of the Czech Academy of Sciences, Zámek 1, CZ-252 43 Průhonice, Czechia

**Keywords:** *Caradmine*, triploid bridge, introgression, speciation, whole genome duplication, polyploidy

## Abstract

- Polyploidisation is a significant reproductive barrier, yet genetic evidence indicates that interploidy admixture is more common than previously thought. Theoretical models and controlled crosses support the ‘triploid bridge’ hypothesis supposing that hybrids of intermediate ploidy facilitate gene flow. However, comprehensive evidence combining experimental and genetic data is missing.
- In this study, we investigated the rates and directions of gene flow within a diploid– autotetraploid contact zone of *Cardamine amara*, a species with abundant natural triploids. We cytotyped over 400 wild individuals, conducted reciprocal interploidy crosses and inferred gene flow based on genome-wide sequencing of 84 individuals.
- Triploids represent a conspicuous entity in mixed-ploidy populations (5%), yet only part of them arose through interploidy hybridisation. Despite being rarely formed, triploid hybrids can backcross with their parental cytotypes, producing viable offspring that are often euploid (in 42% of cases). In correspondence, we found a significant genome-wide signal of gene flow for sympatric, but not allopatric, diploids and tetraploids. Coalescent simulations demonstrated significant bidirectional introgression which is stronger in the direction towards the tetraploid cytotype.
- Triploids, though rare, play a key role in overcoming polyploidy-related reproductive barriers. We present integrative evidence for bidirectional interploidy gene flow mediated by a triploid bridge in natural populations.

## Introduction

Understanding the mechanisms underpinning speciation has been a primary focus of evolutionary biology. The emergence of the biological species concept (Mayr, 1942) underscored reproductive isolation as the ultimate stage of speciation, where intrinsic or extrinsic barriers play pivotal roles in driving the process. Yet, the contribution of intrinsic barriers driven by genetic incompatibilities is still not entirely resolved (Coughlan & Matute, 2020). One of the biological processes acting as an intrinsic barrier to reproduction is whole genome duplication (WGD), which increases the number of sets of chromosomes and is estimated to constitute 15% of speciation events in plants (Wood *et al*., 2009). Unlike other intrinsic barriers, such as Dobzhansky–Muller incompatibilities, which accumulate gradually (Orr & Turelli, 2001; Presgraves, 2010), WGD causes a drastic, immediate change in the genetic composition of an individual within a single generation. The doubling of chromosome sets leads to critical genetic incompatibilities, resulting in developmental and chromosomal imbalances in hybrid progeny resulting from a cross between a newly formed polyploid and its lower-ploidy progenitor. These imbalances are characterised by decreased viability (known as the triploid block) and/or reduced fertility of interploidy hybrids, making whole genome duplication (WGD) a classic example of non-ecological, instant speciation (Otto & Whitton, 2000; Czekanski-Moir & Rundell, 2019).

However, reproductive barriers arising following WGD may still be permeable, as suggested by both theory and experiments, particularly in cases where intermediate cytotypes are viable and could be facilitating backcrossing with plants of parental cytotypes (referred to as the ‘triploid bridge’ pathway – Ramsey & Schemske, 1998; Husband, 2004). In contrast to unidirectional gene flow towards higher ploidy, which occurs when unreduced gametes from a lower ploidy result in hybrids with the ploidy of a higher cytotype (documented e.g. in *Capsella by* Kryvokhyzha *et al*., 2019 and *Betula* by Zohren *et al*., 2016 and Leal *et al*., 2024), a triploid bridge may enable bidirectional gene flow if fertile interploidy hybrids are formed and are capable of backcrossing with both parental ploidies (Kolář *et al*. 2017, Bartolić *et al*., 2024). Indeed, analyses of genetic structure demonstrated interploidy admixture in some allopolyploid (Pinheiro *et al*., 2010; Thórsson *et al*., 2001) and autotetraploid species (Laport *et al*., 2016; Šingliarová *et al*., 2019), but whether this admixture reflects bidirectional gene flow remains untested. Likely ancestral bidirectional gene flow has been documented in *Arabidopsis arenosa* (Arnold *et al*. 2015); however, the virtual lack of natural triploids in mixed-ploidy populations impedes the further interpretation of its mechanistic basis and natural significance (Morgan *et al*. 2020, 2021). Studies investigating interploidy gene flow in nature are often hampered by a lack of genomic tools or resources and by the absence of integration between genomic approaches and experimental tests of pre- and postzygotic barriers. As a consequence, we still lack a satisfactorily described case of bidirectional interploidy gene flow mediated by a triploid bridge mechanism in natural plant populations (reviewed by Bartolić *et al*., 2024). For a more holistic view of introgressive gene flow across a ploidy barrier in a natural environment, we require genetically well-tractable systems with a well-defined cytogeographic distribution and sympatric populations encompassing viable intermediate cytotypes.

One such system is *Cardamine amara* (Brassicaceae), a widely distributed perennial herb inhabiting most of Europe’s wetlands and streams at low to high elevations. In contrast to the diploid cytotype (*C. amara* subsp. *amara*), which is distributed throughout most of Europe, the tetraploid cytotype (*C. amara* subsp. *austriaca*, Marhold, 1999) inhabits the Eastern Alps and neighbouring areas. An overall genetic similarity between diploid and tetraploid cytotypes, together with only slight morphological differentiation and no apparent candidates for interspecific hybridisation, indicates an autopolyploid origin of the tetraploid, likely during the Pleistocene (Lihová *et al*., 2004; Marhold *et al*., 2002; Bohutinská *et al*., 2021). Both cytotypes meet in the northern foothills of the Alps (reaching as far as central Czechia), where they form a distinct secondary contact zone with populations including both diploid and tetraploid individuals as well as several viable individuals of intermediate triploid ploidy growing in proximity (Zozomová-Lihová *et al*., 2015). A well-delimited contact zone, suitable genomic resources and the co-occurrence of both major ploidies together with triploids make *C. amara* an ideal model for addressing the role of the triploid bridge. However, neither the strength of reproductive barriers between diploids and tetraploids nor the intensity of interploidy gene flow and role of triploids therein has been resolved so far.

In this study, we took a multidisciplinary approach involving extensive sampling and cytotyping of plants from mixed-ploidy populations by flow cytometry, reciprocal crossing experiments and population genomics employing whole genome sequencing to answer the following questions: (1) How strong is the triploid block in *C. amara*? (2) Are triploids a product of fusion between a reduced gamete and an unreduced gamete of a diploid or of hybridisation between the major cytotypes? Can triploids mediate interploidy introgression, and if so, do they preferentially produce aneuploid hybrids? (3) Has interploidy introgression left traces in the genome, and if so, has gene flow been stronger towards tetraploids?

## Materials and Methods

### Field sampling

Six populations in different parts of Central Europe were sampled. Two of them were mixed-ploidy populations from the contact zone and four were populations from cytotype-pure areas (Fig. 1a, Table S1). The populations were chosen to include both sympatric populations, where all cytotypes are intermixed, and those deep in cytotype-pure areas away from the contact zone, based on previous thorough flow cytometric screening (Zozomová-Lihová *et al*., 2015). From the cytotype-pure populations, ten individuals were transferred to the greenhouse of Charles University (Prague, CZ) for subsequent crossing experiments and genetic analyses. In the two mixed-ploidy populations (LIP, HLI), multiple geo-referenced individuals were sampled to determine the fine-scale distribution of cytotypes (Fig. 1c, d, e). Following this, ten and ten living individuals of the diploid and tetraploid cytotype and three triploid individuals were transferred from each population for genetic analyses. A subset of these individuals was used in Crossing Experiment 1 (detailed in the Crossing experiments section). Additionally, diploids (2x), triploids (3x) and tetraploids (4x) were sampled later in the LIP population for Crossing Experiment 2, focused on triploid backcrossing. Within each population, individuals were collected with a sampling distance of 3–5 m to minimise the likelihood of collecting individuals resulting from vegetative reproduction. In addition, both populations were revisited during the fruiting season, and seeds were collected from 64 and 96 wild plants of population LIP and population HLI, respectively. The ploidy of these mother plants was unknown at the time of collection and was only determined afterwards, which led to a significant imbalance in the representation of diploid and tetraploid plants in our sample. After confirmation of the ploidy of each mother plant, the collected seeds were germinated, and the ploidy of the seedlings was also determined.

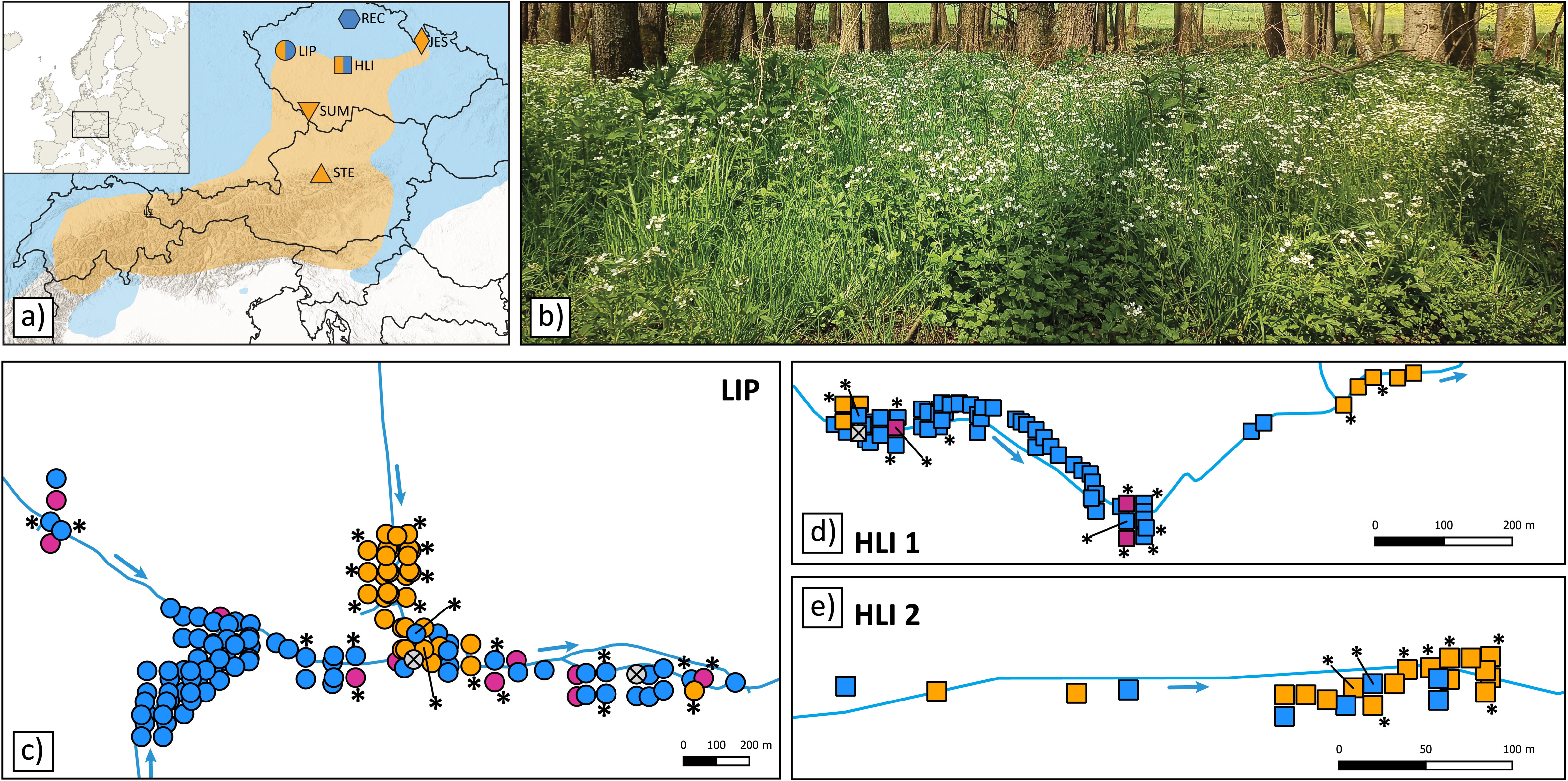

### Ploidy level estimation and chromosome counting

The ploidy level of each individual was estimated separately by measuring relative genome size using flow cytometry with 4,6-diamino-2-phenylindole (DAPI) staining. A two-step protocol, based on Zozomová-Lihová et al. (2015), was followed with *Solanum pseudocapsicum* (2C = 2.59 pg; Temsch et al., 2010) as an internal standard. The prepared nuclei solution was analysed to obtain measurements for a minimum of 3,000 particles using one of two machines, CyFlow ML (Partec) or CytoFlex S (Beckman Coulter), in the Flow Cytometry Laboratory at Charles University in Prague. Relative genome size (RGS), defined as the ratio of the peak fluorescence intensity of the sample to that of the internal standard, was inferred from the resulting histograms using FloMax FCS 2.0 or CytExpert 2.4 software for analyses run on the CyFlow or CytoFlex machines, respectively. Correspondence between the two machines was checked using 20 samples analysed on both instruments.

Individuals were categorised as euploid diploids, triploids or tetraploids based on whether their RGS values fell within the range of three standard deviations above or below the respective means of the established control groups (2x: 0.155–0.191, 3x: 0.262–0.272, 4x: 0.349–0.369). As control groups, we used diploid and tetraploid progeny from the control homoploid crosses from Crossing Experiments 1 and 2. Any values outside these cut-offs were deemed potential aneuploids.

To validate putative aneuploids among individuals with deviating RGS, mitotic chromosome spreads of *C. amara* accessions D7 (2*n* = 16), B6 (2*n* = 32), F8 (2*n* = 31), M8 (2*n* = 31) and L1 (2*n* = 38) were prepared from root tips as described by Mandáková et al. (2019). Chromosome preparations were treated with 100 μg/ml of RNase in 2× sodium saline citrate (SSC; 20× SSC: 3 M sodium chloride, 300 mM trisodium citrate, pH 7.0) for 60 min and with 0.1 mg/ml of pepsin in 0.01 M HCl at 37°C for 5 min; then postfixed in 4% formaldehyde in 2× SSC for 10 min, washed in 2× SSC twice for 5 min, and dehydrated in an ethanol series (70%, 90% and 100%, 2 min each). Chromosomes were counterstained with 4′,6-diamidino-2-phenylindole (DAPI, 2 µg/ml) in Vectashield antifade mounting medium. Fluorescence signals were analysed and photographed using a Zeiss Axioimager epifluorescence microscope and a CoolCube camera (MetaSystems).

### Crossing experiments

Two experiments were performed. In Crossing Experiment 1, diploid plants were crossed with tetraploids to investigate the strength of postzygotic barriers (triploid block; 50 individuals included – 25 diploid and 25 tetraploid Table S2). In Crossing Experiment 2, triploid individuals were crossed with either diploid or tetraploid plants to investigate the potential of natural triploids for backcrossing (triploid bridge; 54 individuals included – 17 diploids, 17 tetraploids and 20 triploids, Table S2). The plants were cultivated in the greenhouse of the Faculty of Science at Charles University in Prague under natural light and with regular watering. When flower buds appeared, flowers designated as pollen acceptors were emasculated to avoid self-pollination. Each emasculated plant was covered with a plastic mesh bag to prevent any unwanted pollination by other plants. After the stigma of an emasculated flower became receptive, we used a mature anther of the designated father plant and rubbed it on the stigma until its surface was fully covered with pollen and then enclosed the pollinated flower in a pollination bag. After pollination, successfully formed siliques were counted and enclosed in paper bags to collect the seeds. In Crossing Experiment 1, we conducted 19 crosses with tetraploids as pollen recipients (4x × 2x; hereinafter, the mother plant is always given first) and 16 with tetraploids as pollen donors (2x × 4x). Additionally, we performed six diploid and ten tetraploid homoploid control crosses (2x × 2x, 4x × 4x). In Crossing Experiment 2, triploids were crossed with diploids and tetraploids in all combinations (2x × 3x [four crosses], 3x × 2x [10 crosses], 3x × 4x [nine crosses], 4x × 3x [five crosses], Table S2) and supplemented them with six diploid and 9 tetraploid control crosses. We aimed to pollinate at least five flowers per cross in both experiments. Seeds obtained from both experiments were counted, weighed using a high-precision analytical scale and germinated in a universal garden substrate under a 21/18°C day/night regimen with 16 hours of light per day. The RGS of each germinated seedling resulting from an interploidy cross and a subset of 40 (20 diploid and 20 tetraploid) seedlings from control crosses was measured by flow cytometry as described above.

For the statistical evaluation of each experiment, two models were established and compared. Initially, a null model assumed a constant mean value for the dependent variable (number of seeds per silique, average mass per seed, proportion of germinated seedlings) across all individuals. Subsequently, a more complex model allowed the mean value to vary among different cross-types. The significance of the cross-type effect was assessed by means of likelihood ratio tests comparing the models. Linear models, calculated using the R package lme4 (Bates *et al*., 2015), were used to determine relative differences in seed set and seed mass, and Tukey post-hoc tests were then employed to compare individual treatments. Differences in germination rate were calculated using generalised linear models with a binomial distribution of residual variation in the R package lme4 (Bates *et al*., 2015). The proportion of germinated seedlings per seed parent was used as a dependent variable, and the type of cross was used as a predictor.

### DNA extraction, library preparation, sequencing, and raw data processing and filtration

Eighty-four individuals from six populations were selected for population genomic studies: 39 from cytotype-pure populations and 45 individuals from mixed-ploidy populations (for details see Table S3). Genomic DNA was extracted from silica-gel-dried leaves using the sorbitol extraction method and then purified using AMPure XP (Beckman Coulter Inc., Brea, California, USA). Samples were genotyped for genome-wide single nucleotide polymorphisms (SNPs) using the whole genome sequencing protocol LITE of Perez-Sepulveda et al. 2021. The libraries were sequenced in 300 cycles (2 × 150 bp paired-end/PE reads) on the Illumina NovaSeq platform.

Illumina sequencing reads were demultiplexed using the fastx_barcode_splitter.pl script from the FASTX-Toolkit v. 0.0.14. Read ends and reads where the average quality within the 5-bp window fell below Q20 were trimmed, and reads of less than 50 bp were discarded by Trimmomatic v. 0.36 (Bolger *et al*., 2014). The resulting reads were compressed into clumps, and duplicates were removed using the script clumpify.sh (BBTools; https://jgi.doe.gov/data-and-tools/bbtools).

The sequencing reads were mapped on to a reference genome of *C. amara* (Bohutínská *et al*. 2021) using BWA 0.7.3a (Li, 2013) and the resulting BAM files were processed the Picard Toolkit v. 2.22.1. Variant calling was performed for each individual using the HaplotypeCaller module from the Genome Analysis Toolkit v. 3.7-0 (GATK; McKenna *et al*., 2010), specifying the ploidy of each individual. Next, variants were aggregated and genotyping across all individuals was performed using the GATK’s GenotypeGVCFs module, which is suitable for joint genotyping of mixed-ploidy datasets (see e.g. Monnahan *et al*. 2019). Variant filtration was performed by the VariantFiltration module, requiring a minimum sequencing depth of 8x and applying the filter parameters indicated by GATK’s best practices (Van der Auwera *et al*., 2013). Finally, the SelectVariants module was used to capture putatively neutral 4-fold degenerated biallelic sites, passed filter parameters, with no more than 20% of missing genotypes, but excluding genes that showed excess heterozygosity or read depth (potential paralogues mapped on top of each other). Four-fold degenerate SNPs were identified by the Identify_4D_Sites.pl script (available at https://github.com/tsackton/linked-selection). Genes with excess heterozygosity (fixed heterozygous in at least 3% of SNPs in one or more diploid populations) and sites with read depth exceeding the mean plus double the standard deviation with at least 10% of samples were identified following Šlenker (2022).

### Population genomic analyses

Initially, genetic clusters were inferred using the Bayesian clustering algorithm implemented in STRUCTURE v. 2.3.4 (Pritchard *et al*., 2000), giving unbiased results with mixed-ploidy populations (Stift *et al*. 2019). STRUCTURE analyses required unlinked SNPs, which were obtained by randomly selecting a single SNP from 20,000-bp windows using the vcf_prune.py script (Šlenker, 2024). To capture the overall data variability, 100 datasets with randomly selected SNPs were analysed. Each dataset was analysed for each K = 1–7, with a burn-in length of 100,000 generations and data collection for an additional 900,000 generations, setting the admixture model and correlated allele frequencies. The results for 100 datasets were averaged using the programme CLUMPP (Jakobsson & Rosenberg, 2007) and drawn with the DISTRUCT routine (Rosenberg, 2004). The approach of Evanno *et al*. (2005) was adopted to determine the optimal K value. Secondly, we displayed genetic distances among individuals using principal component analysis (PCA) based on Euclidean distance as implemented in adegenet v1.4-2 (Jombart, 2008). Thirdly, we calculated Nei’s (Nei, 1972) distances among all individuals using the StAMPP package (Pembleton *et al*., 2013), developed specifically for analysing SNP datasets on mixed-ploidy scenarios, and displayed them using the neighbour network algorithm in SplitsTree (Huson & Bryant, 2006).

### ABBA–BABA test and demographic modelling

To quantify the extent of recent introgression in the sympatric populations, we conducted an ABBA–BABA test, which relies on Patterson’s D statistic to estimate the genome-wide excess of shared derived alleles between two taxa (Green *et al*., 2010; Martin *et al*., 2015). This test assumes that both the ABBA and BABA topologies occur at the same frequency in accordance with the incomplete lineage sorting (ILS) hypothesis. However, if there is introgression between two taxa of a bifurcating tree, one topology occurs with much greater frequency than the other. One allopatric tetraploid population far from the contact zone (STE) was used as P1 in all cases. For sympatric combinations, tetraploid and diploid individuals from each mixed population (LIP, HLI) were used as P2 and P3, respectively. For allopatric combinations, each tetraploid population occurring outside the contact zone (SUM, JES) was used as P2, and the only allopatric diploid population (REC) was used as P3. One population of the closely related but geographically distant taxon *C. amara* subsp. *balcanica* was used as an outgroup. To calculate the D statistic, we used scripts written by Simon Martin available at https://github.com/simonhmartin/tutorials/tree/master/ABBA_BABA_whole_genome.

To further test for the presence of gene flow between diploids and tetraploids and to quantify its potential asymmetry, we used fastsimcoal v. 2.709 (Excoffier *et al*., 2021). For a pair of diploid and tetraploid (sub-)populations from each mixed-ploidy population (LIP and HLI) separately, we constructed folded two-dimensional site frequency spectra (SFS) from the variant and invariant four-fold degenerated sites (filtered in the same ways as above) using python scripts FSC2input.py available at https://github.com/pmonnahan/ScanTools/ (Monahan *et al*., 2019). For each population pair, we compared the following four scenarios (Fig. S1): (1) no gene flow or migration, (2) unidirectional gene flow from diploids to tetraploids, (3) unidirectional gene flow from tetraploids to diploids, and (4) equal bidirectional gene flow between diploids and tetraploids. For each scenario and population pair, 50 fastsimcoal runs were performed. For each run, 40 ECM optimisation cycles were allowed to estimate the parameters, and 100,000 simulations were conducted at each step to estimate the expected SFS. Further, the partition with highest likelihood for each fastsimcoal run was identified and values of the Akaike information criterion (AIC) for these partitions were calculated and summarised across the 50 fastsimcoal runs. The scenario with the lowest median AIC value within each population was considered the most favourable.

## Results

### Cytotype structure in the contact zone and within mixed-ploidy populations

Three main ploidy levels with RGS values corresponding to diploids, triploids and tetraploids were detected among the 356 field-collected adult individuals (Table S1). In addition, three putative aneuploids with RGS values in between triploid and diploid (two plants) and between triploid and tetraploid (one plant) were sampled in the mixed-ploidy populations LIP and HLI, respectively (Fig. 1c, d, e; Table S4). The fine-scale distribution of cytotypes in the two mixed-ploidy populations revealed the presence of both ploidy-uniform patches and parts where all three cytotypes grow only a few metres apart (Fig. 1c,d,e). The seeds collected in natural mixed-ploidy populations had variable but generally meagre germination rates (0–60%, 8% on average, Fig. S2). Among 122 germinated seedlings, the RGS values corresponded to both euploid (84%) and aneuploid (16%) values (Fig. S3). Seedlings of both diploid and triploid mothers exhibited values corresponding to the diploid state (79% and 21% of seedlings from diploid and triploid mothers, respectively). The rest were putative aneuploids with RGS values between diploids and tetraploids, with the exception of one putative tetraploid found among the progeny of a diploid parent. The RGS of seedlings from tetraploid mothers ranged from values corresponding to hypo-tetraploid (15%) and tetraploid (62%) up to hyper-tetraploid (23%) aneuploid (Fig. S3). No seedling with RGS corresponding to the triploid state was found.

### Strength of the interploidy barrier inferred from crossing experiments

We first crossed diploid and tetraploid plants to investigate the strength of postzygotic barriers (triploid block, Crossing Experiment 1). Difference in the seed set was significant overall (F_3, 47_ = 3.71, p = 0.018, Fig. S4a), yet rather small, as it was only the category of homoploid control 4x × 4x crosses that was significantly higher than one of the interploidy crosses (2x × 4x; Fig. S4a). The seed mass was also different (*F*_3, 47_ = 98.62, p < 0.001), being similar between the interploidy crosses but markedly lower than that of homoploid crosses of both types (Fig. S4b). Consequently, there was also significant variation in germination rates among the crossing treatments (Fig. 2a, χ^2^ = −339.2, *df* = 3, *p* < 0.001). The progeny of homoploid controls had higher germination percentages (40% and 55% for diploid and tetraploid crosses, respectively) than interploidy crosses, in which no viable seeds were produced by diploid seed parents and only a single germinable seed (< 0.5%) was formed after pollination of a tetraploid by a diploid pollen donor. The RGS value of this plant corresponded to a triploid (Fig. 2a).

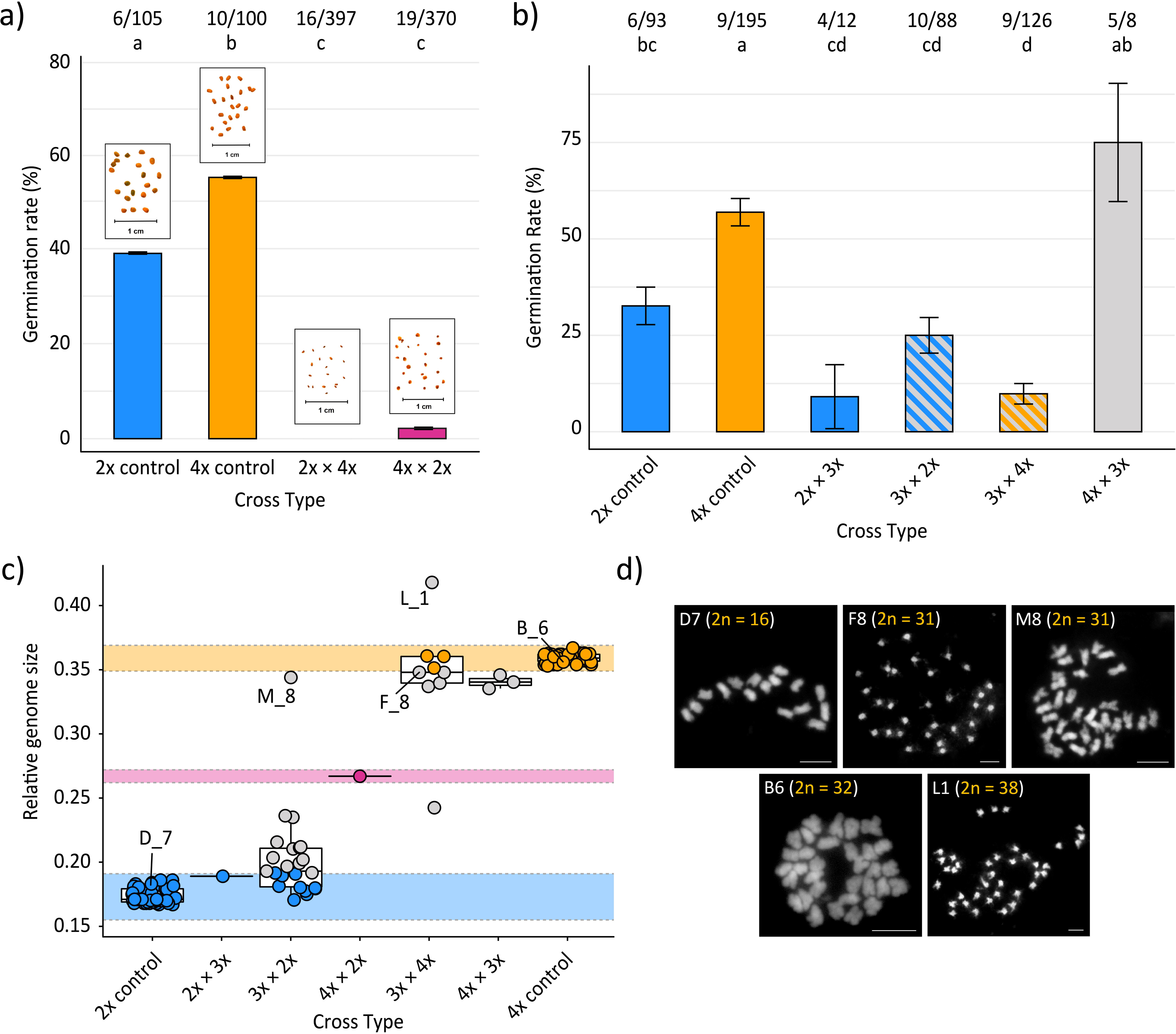

Then, triploid plants were crossed with diploids and tetraploids to investigate the potential of natural triploids to act as introgression mediators (i.e. as part of the triploid bridge pathway, Crossing Experiment 2). There was a significant difference in seed set between successful crosses (*F*_5, 58_ = 5.06, *p* < 0.001), with larger seed sets in tetraploid control crosses than in the majority of interploidy crosses (Fig. S4c). There was no significant difference between the diploid control and the interploidy crosses, with average seed set values of interploidy crosses ranging from 0.3 to 5.5 seeds per silique. There was no statistically significant difference in seed mass among the different types of crosses (*F*_5, 58_ = 0.94, *p* = 0.4608; Fig. S4d). The germination rates, again, differed significantly between the different types of crosses (χ^2^ = 97.018, *df* = 5, *p* < 0.001, Fig. 2b), although the difference was not as pronounced as in the previous experiment. Tetraploid controls generally had significantly higher germination than the other types of crosses, with the exception of one type of backcross that was similar (4x × 3x). The germination rates of triploid backcrosses were highly variable but always non-zero (8–75% across treatments; Fig. 2b).

The relative genome size of the plants obtained from successful triploid backcrosses (36 cytotyped plants) corresponded to both euploid and aneuploid values defined based on the relative deviation of RGS from the control euploid values. The ploidy of the crossing partner of a triploid (diploid or tetraploid) significantly affected the proportion of putatively euploid progeny (χ² = 21.78, *df* = 1, *p* < 0.001). Crosses of triploids with diploids resulted in a higher proportion of putatively euploid progeny (41%, with RGS values corresponding to diploids) than crosses involving tetraploids, where the proportion of euploid (tetraploid) progeny was 25% (Fig. 2c). Chromosome counts obtained from three individuals classified by RGS as aneuploids confirmed the aneuploid number of chromosomes: F_8 (3x × 4x) – hypotetraploid, 2n = 31; L_1 (3x × 4x) – hypopentaploid, 2n = 38; M_8 (3x × 2x) – hypotetraploid, 2*n* = 31 (Fig. 2d).

### Genetic structure based on genome-wide SNPs

Sequencing of 84 individuals from six populations (Fig. 3a) produced between 24,453,671 and 84,627,579 reads per sample, averaging 38,264,942.5 reads after quality control and deduplication (Table S3). Of these, 37% to 94.2% were successfully mapped onto the reference genome, with an average mapping rate of 83.55%. The final VCF file contained 1,448,166 filtered putatively neutral four-fold degenerate SNPs with an average depth of coverage of > 30× that were used in subsequent analyses. Bayesian clustering in STRUCTURE, based on a subset of 7,169 LD-pruned SNPs, suggested K = 2, 3 and 4 as stable partitions (strong similarity across runs), with K = 3 exhibiting the highest relative likelihood difference (delta K). Diploids and tetraploids from cytotype-pure populations separated already under K = 2 (Fig. 3d). Additional separation of tetraploid individuals from mixed-ploidy population LIP was observed under K=3 (Fig. 3d) and K= 4 (population JES, Fig. S5). Tetraploids from the two mixed-ploidy populations, HLI and LIP, however, showed a high proportion of diploid cluster ancestry under K = 2. The separation of pure diploid and tetraploid populations, in contrast to the closer position of both major cytotypes from mixed-ploidy populations, was also supported by the neighbour-joining network (Fig. 3b).

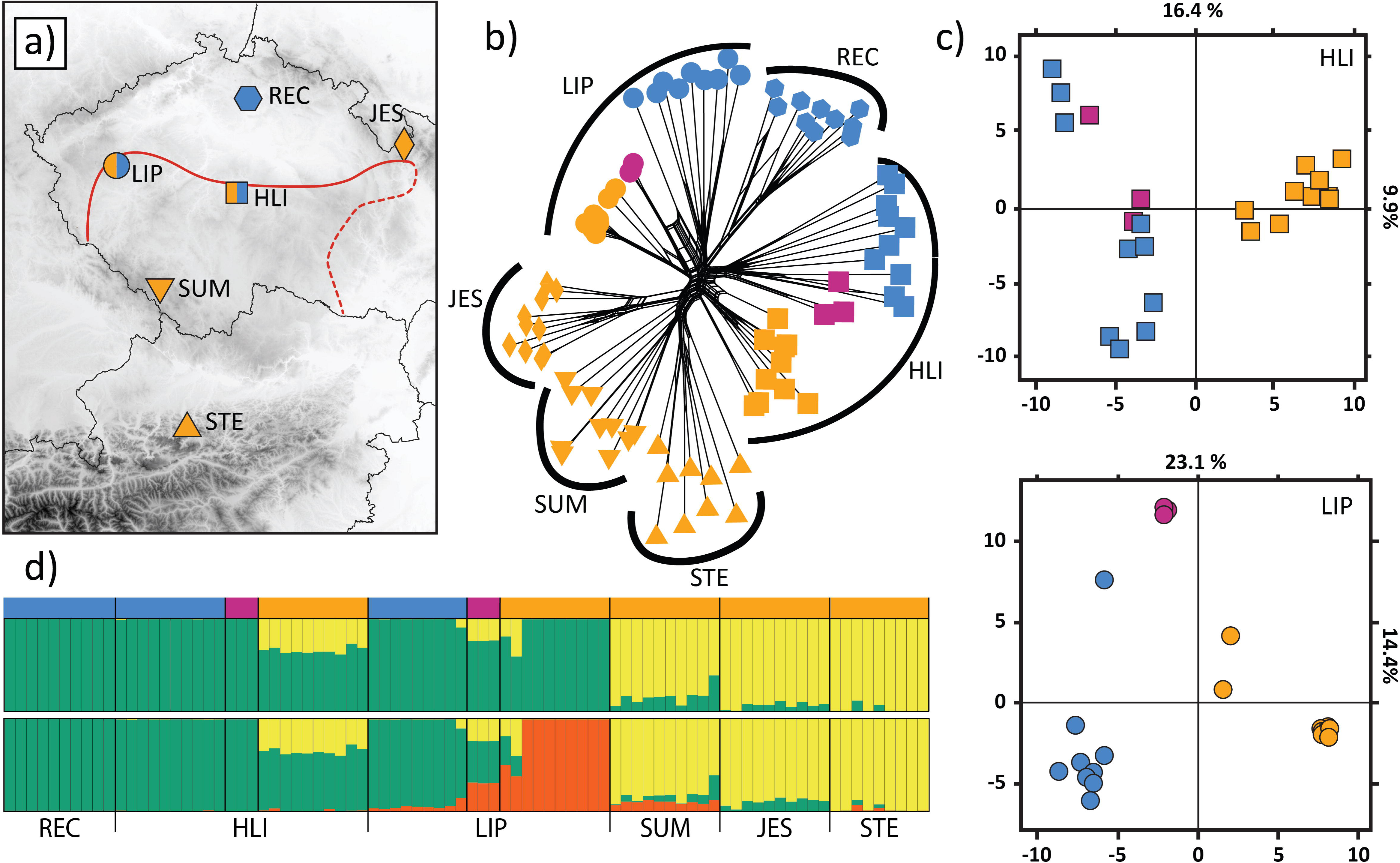

Interestingly, triploids from both mixed-ploidy populations showed contrasting assignment patterns. Triploids from population HLI clustered exclusively with their sympatric diploids in both STRUCTURE and neighbour-joining network analysis whereas triploids from population LIP were a mixture of both STRUCTURE clusters and occupied intermediate positions in the network (Fig. 3b, d). The contrasting genetic make-up of triploids was further corroborated by principal component analyses run separately for each mixed-ploidy population where HLI triploids clustered with diploid individuals; however, LIP triploids occupied an intermediate position between diploids and tetraploids (Fig. 3c).

### Interploidy introgression and the direction of gene flow

We tested for the presence of interploidy introgression using a four-taxon test (ABBA–BABA) by setting different combinations of allopatric populations differing in ploidy and sympatric diploid and tetraploid sub-populations as donors/recipients of introgression and spatially distinct tetraploid populations outside the contact zone (STE) as P1 (Fig. 4a). The analyses revealed significant interploidy introgression in mixed-ploidy populations whereas no significant admixture was observed between tetraploids and diploids sampled outside the contact zone. Tree topologies testing for introgression between tetraploid and diploid individuals from mixed-ploidy populations LIP and HLI resulted in significant D values of 0.28 and 0.36, respectively (Table S5). On the contrary, low and statistically non-significant D values were found for tree topologies involving tetraploid (SUM, JES) and diploid (REC) populations further away from the contact zone, demonstrating a lack of detectable admixture in pure-ploidy allopatric populations (Fig. 4a).

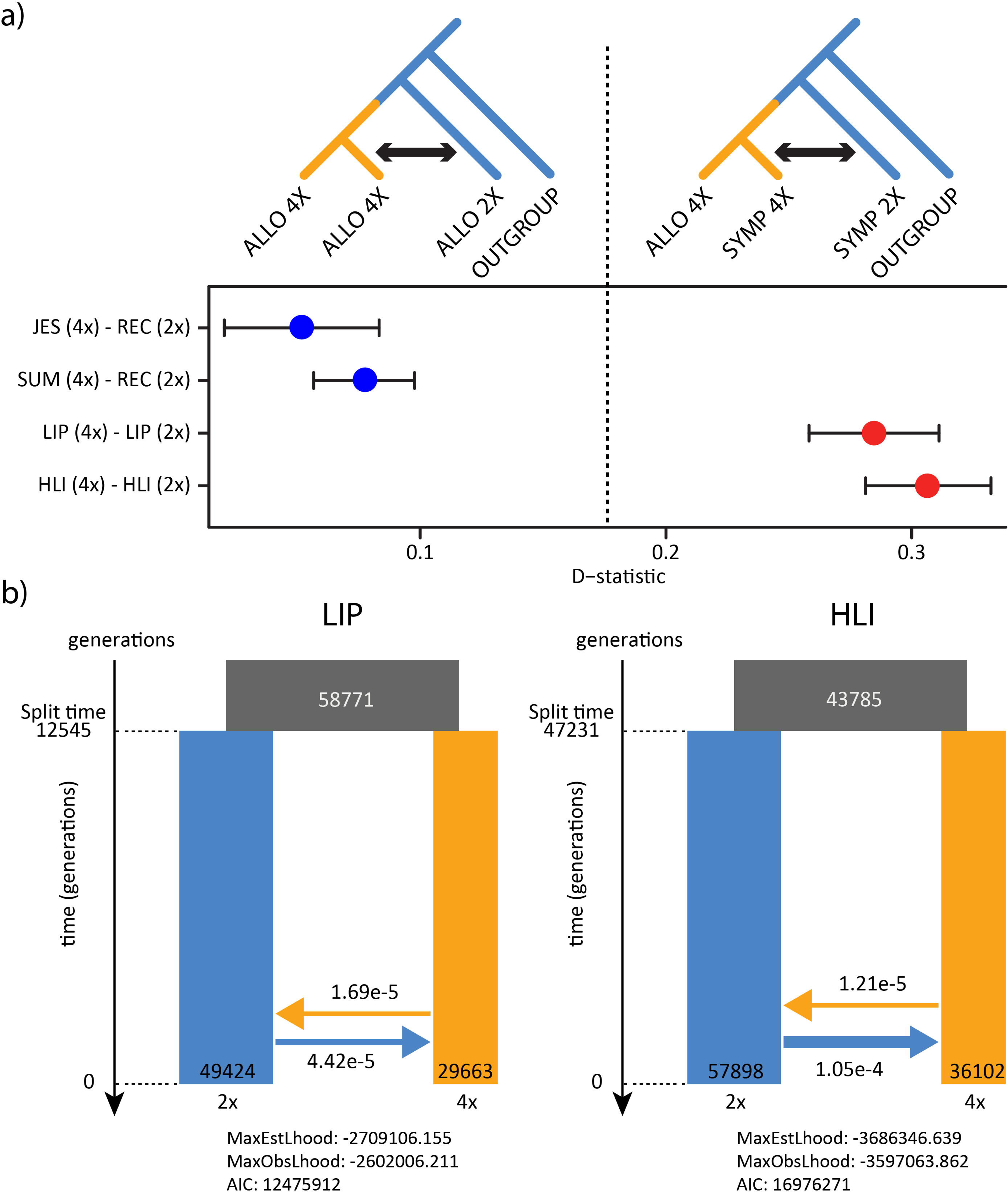

To complement the introgression tests, the strength and direction of gene flow in the two mixed-ploidy populations were also estimated using coalescent simulations. The scenario assuming bidirectional interploidy gene flow exhibited the lowest median and absolute AIC values with both populations (Fig. 4b). The second-best scenario, assuming only unidirectional 2x-to-4x (forward in time) gene flow, was markedly worse (ΔAIC = 3,913.45) than the bidirectional scenario in both the LIP and HLI populations (median ΔAIC = 3,022.81 and 4,439.32, respectively; Fig. S6). The estimated migration rate (i.e. the probability of an individual sprouting in one ploidy subpopulation from a seed originating in another one over the course of one generation) was greater in the direction from diploids to tetraploids, forward in time (4.42 × 10^−5^ and 1.05 × 10^−4^ for population LIP and population HLI, respectively), compared to gene flow from tetraploids to diploids (1.69 × 10^−5^ and 1.21 × 10^−5^ for population LIP and population HLI, respectively).

## Discussion

In this study, we explore the pathways and rates of bidirectional interploidy gene flow in natural populations of a mixed-ploidy plant species. By innovatively combining field surveys, crossing experiments and population genomics, we present robust evidence of significant gene flow between different ploidy levels across multiple natural populations. Specifically, we describe the pathway by which triploid hybrids form in *C. amara* and document their persistence in nature and their ability to backcross. Furthermore, we demonstrate genome-wide signals of bidirectional introgression between diploid and tetraploid individuals growing in natural sympatry. In the following subsections, we discuss the mechanisms and rates of triploid formation and further implications of triploid bridges for plant evolution through whole genome duplication.

### A strong but incomplete postzygotic barrier to triploid formation

Our crossing experiments between diploids and tetraploids have revealed a very strong post-pollination barrier, evidenced by the fact that only one viable triploid plant was produced. A triploid block, decreased hybrid seed viability caused by the parent-of-origin epigenetic imbalance in endosperm development (Köhler *et al*., 2010, 2021), is the likely underlying mechanism. This is suggested by the formation of large numbers of low-germinable and overall malformed seeds in experimental interploidy crosses but not in controls. Similar phenotypes have been found in other Brassicaceae species, for example, of the genera *Brassica* and *Arabidopsis*, for which this mechanism has been comprehensively documented (Scott *et al*., 1998; Stoute *et al*., 2012; Morgan *et al*., 2021). In addition, the viable triploid seed was formed in a cross when tetraploid acted as a mother plant, i.e. the cross direction that usually produces more viable offspring also in other plants with triploid block (Morgan *et al*., 2021, Bartolić *et al.,* 2024). However, in natural *C. amara* populations, triploids constitute a significant, persistent entity, accounting for 5–6% of adult individuals within mixed-ploidy populations, consistently occurring across multiple years in the same spots and for over fifteen years in the same populations (Krásná, 2008; Zozomová-Lihová *et al*., 2015). This disparity between the outcomes of experimental and naturally occurring crosses contradicts typical observations. In several other plant polyploid systems, experimental crosses have shown a ‘leaky’ triploid block, yet hybrids were scarce or non-existent in natural populations (Greiner & Oberprieler, 2012; Sonnleitner *et al*., 2013; Hülber *et al*., 2015; Morgan *et al*., 2020, 2021; Šemberová *et al*., 2021). This discrepancy has been attributed to factors such as prezygotic barriers, reduced hybrid fitness, or a combination of both. In the case of *C. amara*, there is no indication of strong prezygotic barriers, such as temporal isolation or pollinator preference, as all cytotypes are morphologically indistinguishable (Marhold, 1999), coexist in immediate proximity and overlap in flowering phenology (personal observation).

Considering the strong triploid block, the relatively frequent presence of triploids in nature can most likely be attributed to the life history of the species, particularly its perenniality and clonal reproductive strategy. In the case of *C. amara*, vegetative reproduction is frequent and vital (Koch *et al*., 2003), which may enhance the longevity and persistence of triploid hybrids once they have formed. Previous studies have shown that polyploids often depend on vegetative reproduction, which not only safeguards many nascent polyploids from extinction, but also plays a crucial role in polyploid speciation by facilitating more efficient space utilisation and decreasing mortality from small-scale disturbance events (Herben et al., 2017; Van Drunen & Husband, 2019; Van Drunen & Friedman, 2022).

### Triploids as mediators of interploidy gene flow

In spite of the traditional assumption that polyploidisation is an instantaneous and perfect barrier, steeply accumulating genomic evidence documents that interploidy gene flow is frequent and forces the re-appraisal of its relevance for the formation, establishment and further evolution of novel polyploid lineages (Schmickl & Yant, 2021; Bartolić *et al*., 2024; Brown *et al*., 2024). Even though there are multiple scenarios of interploidy admixture in which introgression is primarily unidirectional from diploids to polyploids, triploids serve as an essential conduit for gene flow in the reverse direction, enabling also introgression from polyploids to diploids (Petit *et al*., 1999; Kolář *et al*., 2017; Bartolić *et al*., 2024). Triploids are present in over 60% of well-established mixed-ploidy systems comprising diploids and tetraploids (Kolář *et al*., 2017), yet detailed genetic studies on their role in extant gene flow are limited because researchers have primarily focused on the role of triploids in the formation of new polyploids (Bretagnolle & Thompson, 1995; Ramsey & Schemske, 1998; Husband, 2004). Triploid formation entails either a diploid–tetraploid cross (van Dijk & van Delden, 1990; Peckert & Chrtek, 2006; De Hert *et al*., 2012; Sabara *et al*., 2013; Vallejo-Marín *et al*., 2016; Popelka *et al*., 2019; Castro *et al*., 2020) or the fusion of one reduced and one unreduced diploid gamete (Slovák *et al*., 2009; Schinkel *et al*., 2017; Šmíd *et al*., 2020). These two mechanisms have only rarely been found to coexist in the same system by studies based on cytotype distribution patterns (Mandák *et al*., 2016); however, sufficient sampling combined with thorough genotyping may reveal that such a pattern is more frequent. Here we present genetic evidence for both pathways: diploid-tetraploid hybridisation in population LIP and fusion of reduced and unreduced diploid gametes in population HLI (Fig. 3, Fig. 4). Such a result also implies that estimating the levels of interploidy gene flow solely based on triploid frequency may be misleading, as even triploids found in mixed-ploidy populations may not always be hybrids (Bartolić *et al*., 2024).

In addition to triploid formation, we also show that triploid hybrids are fertile and capable of backcrossing both in experimental and natural conditions. These results add to a body of evidence primarily based on pollen fertility assessments (Ramsey & Schemske, 1998; Laport *et al*., 2016; Morgan *et al*., 2021) that (partially) fertile triploid hybrids may further contribute to the composition and dynamics of the contact zones, lending support to theoretical models (Husband, 2004; Kauai *et al*. 2024). In a novel finding, we also show that the relative genome size of a significant (42%) proportion of triploid backcross progeny corresponds to either the diploid or the tetraploid level, demonstrating the potential of triploids as mediators of interploidy introgression between the two major euploid cytotypes, in line with genomic data (see the following subsection). Notably, however, the majority of progeny resulting from triploid backcrosses were still aneuploids, and some additional aneuploids differing by a single chromosome might have been misclassified as euploids because of the limited resolution of our flow cytometric approach. On the other hand, aneuploids might also play a role in mediating interploidy gene flow in experimental populations of *Arabidopsis thaliana* (Henry *et al*., 2005, 2009). Interestingly, we also detected three viable adult individuals in the mixed-ploidy populations, with RGS corresponding to aneuploid values suggesting that aneuploids may form and survive until adulthood also in natural populations, similarly as has been observed in contact zones between cytotypes of *Tripleurospermum inodorum* (Čertner *et al.,* 2017).

### Bidirectional and asymmetric interploidy gene flow

In both mixed-ploidy populations, the coalescent models supported ongoing bidirectional gene flow, aligning with both experimental findings and field observations. This also supports the involvement of triploid individuals as mediators of gene flow, as there is no alternative mechanism by which introgression could proceed from tetraploids towards diploids (Bartolić *et al.,* 2024). Moreover, differences in the intensity of gene flow between the two investigated populations align with the distribution and frequency of cytotypes in the field. Gene flow towards diploids is more pronounced in population LIP, which exhibits a more intermingled, mosaic-like cytotype structure and also harbours triploid individuals that have been proven to be hybrids. By contrast, signals of gene flow towards diploids are weaker in population HLI, where triploids are currently segregated from tetraploids and occur within diploid patches with a genetic profile close to diploids. This suggests that triploids might play a significant role in population LIP by mediating gene flow to diploids whereas in population HLI we have not found any conclusive evidence for any ongoing interploidy hybridization via triploids, at least based on our current sampling. Although bidirectional interploidy gene flow has been expected based on theoretical models (Husband, 2004; Kauai *et al*., 2024; Felber & Bever, 1997), its presence has been suggested only rarely in natural systems, mostly based on indirect evidence of genetic clusters spanning cytotypes (Ståhlberg, 2009; Nierbauer *et al*., 2017; Šmíd *et al.,* 2020). Our data, in a testable framework based on coalescent simulations, provide evidence for bidirectional gene flow. We speculate that gene flow in *C. amara* is ongoing (population LIP) or at least recent (population HLI), reflecting the presence of fertile triploids in the field and genetic support for the presence of introgression in mixed-ploidy populations but not outside the contact zone.

In both mixed-ploidy populations, gene flow was inferred to be stronger in the direction towards tetraploids. Such an asymmetry likely reflects an additional route of unidirectional gene flow from lower to higher ploidy: the merger of an unreduced gamete of a diploid with a reduced gamete of a tetraploid leading to hybrid tetraploid progeny. Indeed, previous extensive crossing experiments often found a certain fraction of such tetraploid hybrids demonstrating that this pathway may act in addition to a triploid bridge (e.g. van Dijk & van Delden, 1990; Burton and Husband 2001; Chrtek *et al*., 2017; Sutherland & Galloway, 2017; Castro 2020; Morgan *et al*., 2021). Although we did not encounter such a hybrid in our limited crossing experiment, the observation of a tetraploid seedling among the progeny of a diploid seed parent sampled in population HLI demonstrates that unreduced gametes of diploids may also be involved in tetraploid formation in the field. The overall importance of this pathway is illustrated by the fact that in most well-documented cases of interploidy gene flow, the direction is typically inferred as unidirectional, from lower to higher ploidy levels (Zohren *et al*., 2016; Kryvokhyzha *et al*., 2019; Monnahan *et al*., 2019; Wang *et al*., 2023; Leal *et al*., 2024; Ding *et al*., 2024). Considering that unreduced gametes are present in many systems (Ramsey & Schemske, 1998; Kreiner *et al*., 2017a, b), it is no surprise that this direction is more prevalent in nature (Stebbins, 1971; Brown *et al*., 2024; Bartolić *et al*., 2024). In sum, our field, experimental and genomic data provide a coherent picture of ongoing interploidy gene flow at multiple sites of the interploidy contact zone. The hybridisation is likely bidirectional yet asymmetrically stronger in the direction towards tetraploids, suggesting the involvement of both triploid hybrids and unreduced gametes.

Interploidy gene flow may be an important mechanism of how the gene pool of a nascent, initially depauperate polyploid may be enriched. Specifically, in cases involving autotetraploids, recent genomic evidence suggests that such gene flow provides novel genetic variation that can benefit polyploid adaptation (Baduel *et al*., 2018, reviewed in Schmickl & Yant, 2021). Interploidy gene flow also constitutes a key component of a pathway enabling gene flow between different species. Mathematical models have shown that WGD-mediated gene flow can serve as a bridge between diploid lineages, where introgression is otherwise impeded by interspecies barriers (Kauai *et al*., 2024). This lateral transfer of genes from polyploids to diploids has been observed in several grass species, particularly with genes coding for the C4 photosynthetic pathway (Christin *et al*., 2012; Phansopa *et al*., 2020). The spread of genetic variation to the diploid level is facilitated exclusively by triploid crosses, highlighting the crucial role of the triploid bridge in intra- and interspecific genetic exchange and possibly adaptation.

## Conclusion

The triploid bridge is a fundamental concept in polyploid biology, offering a mechanistic explanation for bidirectional gene flow in polyploid complexes. Our study not only documents this pathway, but also demonstrates its genomic impact in natural plant populations, underscoring the critical role of triploid hybrids as mediators of gene flow. On the other hand, our data also reveal that not all triploid individuals in mixed-ploidy populations are hybrids, and alternative pathways for their origin via autopolyploidzation from diploids may be involved. Remarkably, despite the strong triploid block that limits their formation, we observed a significant frequency of triploid hybrids in natural populations. This likely reflects the capacity of *C. amara* to spread and persist clonally, allowing triploids to maintain their presence over extended periods. Our findings validate theoretical predictions and demonstrate that odd-ploidy cytotypes, though rare and often of reduced fertility, can significantly influence the genomic landscape of natural populations. We anticipate that future multi-species studies will uncover the broader prevalence of triploid bridges across diverse species and clarify the role of species traits, such as clonality, in facilitating substantial bidirectional interploidy gene flow.

## Acknowledgements

This work was supported by the Czech Science Foundation (project Nos 23-07204M led by F.K. and 22-16826S led by T.M.), the Grant Agency of Charles University (GAUK, project No. 383621 led by P.B.), and the Slovak Research and Development Agency (project no. APVV-21-0044 led by K.M.). Additional support was provided by the Czech Academy of Sciences (long-term research development project No. RVO 67985939). The sequencing was performed by the Norwegian Sequencing Centre at the University of Oslo. Computational resources were provided by the e-INFRA CZ project (ID:90254), supported by the Ministry of Education, Youth and Sports of the Czech Republic.

## Data availability

The raw resequencing data have been deposited in the BioProject collection of biological data, maintained by the National Center for Biotechnology Information (NCBI), under ID PRJNA1148746 and SRA accession Nos SRR30265016–SRR30265099.

## Supporting information

Additional supporting information may be found in the online version of this article.

**Fig. S1** Cartoons illustrating four demographic scenarios for current gene flow between diploids and tetraploids in mixed-ploidy populations.

**Fig. S2** Germination rate of field-collected seeds of *C. amara* from the localities Lipí and Hliňánky from 2020, 2021 and 2022.

**Fig. S3** Relative genome size of germinated seeds collected from mixed-ploidy populations HLI and LIP.

**Fig. S4** Performance of progeny from Crossing Experiment 1 (left column; a, c) and Crossing Experiment 2 (right column; b, d) assessed based on selected fitness traits measured across different types of crosses obtained by hand pollination of *C. amara*.

**Fig. S5** Population structure, returned by STRUCTURE at K = 2 to K = 6, of six populations of *C. amara* in Central Europe.

**Fig. S6** Comparison of Akaike information criterion (AIC) values across four scenarios approximating the type of gene flow between diploids and tetraploids in mixed-ploidy populations.

**Table S1** Sampling details of populations of *C amara* and average relative genome size of individuals from these populations.

**Table S2** Summary of crossing experiments.

**Table S3** Details on the 84 sequenced individuals.

**Table S4** Relative genome sizes of putative aneuploid individuals from field samples and experimental progeny with chromosome counts.

**Table S5** Results of ABBA-BABA tests for introgression detection.

