## Supplemental_Figures for "Overcoming ploidy barriers: the role of triploid bridges in the genetic introgression of *Cardamine amara*"

### *New Phytologist* Supporting Information

**Article title: Triploid bridge mediates introgression across a ploidy barrier in the natural contact zone of *Cardamine amara***

Authors: Paolo Bartolić, Alena Voltrová, Lenka Macková, Gabriela Šrámková, Marek Šlenker, Terezie Mandáková, Nelida Padilla García, Karol Marhold, Filip Kolář

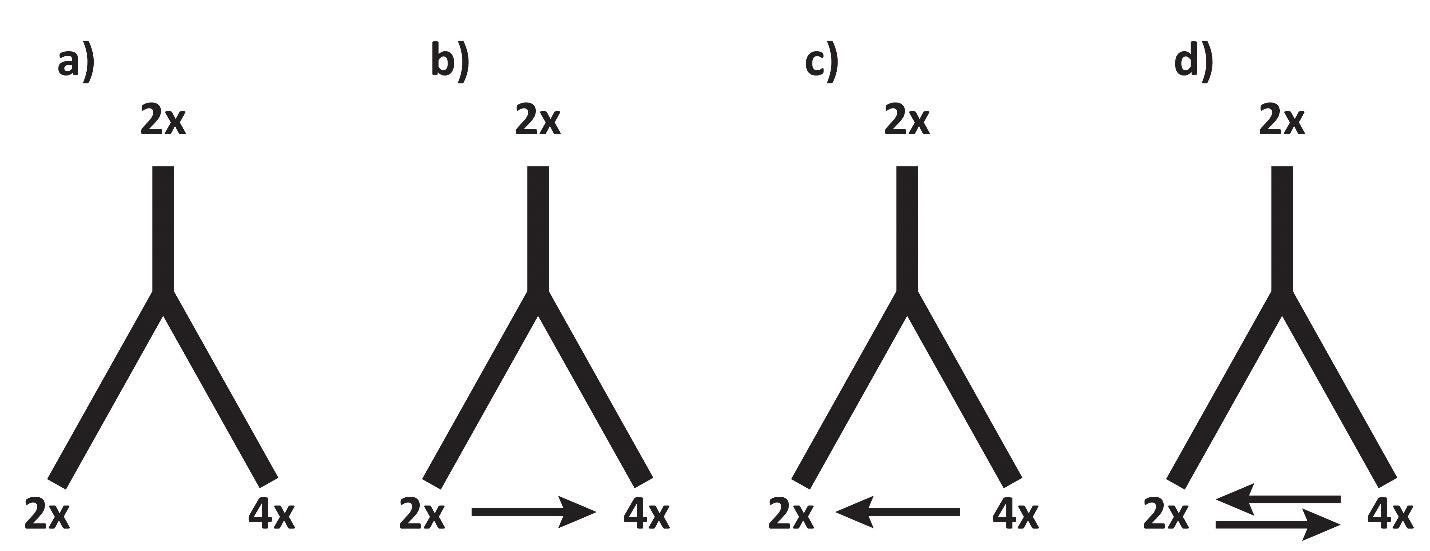

**Fig. S1** Cartoons illustrating four demographic scenarios for current gene flow between diploids and tetraploids in mixed ploidy populations addressed using fastsimcoal. a) no gene flow (migration), b) unidirectional gene flow from diploids to tetraploids, c) unidirectional gene flow from tetraploids to diploids, and d) equal bidirectional gene flow between diploids and tetraploids.

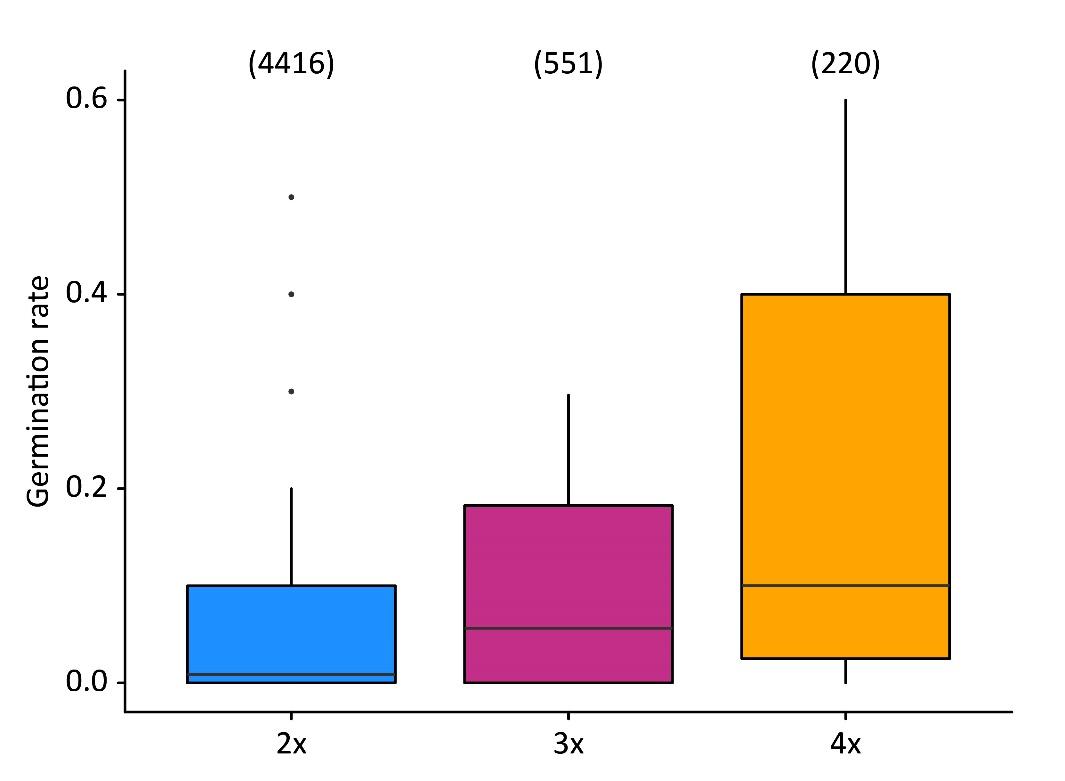

**Fig. S2** Germination rate of seeds of *C. amara* sampled in the field from mothers from the mixed-ploidy localities Lipí and Hliňánky. Numbers of seeds sown are above boxplots; the different sample sizes reflect different success in finding fertile individuals with developed ripe seeds at the time of collection (multiple collections in years 2020-2022 were done).

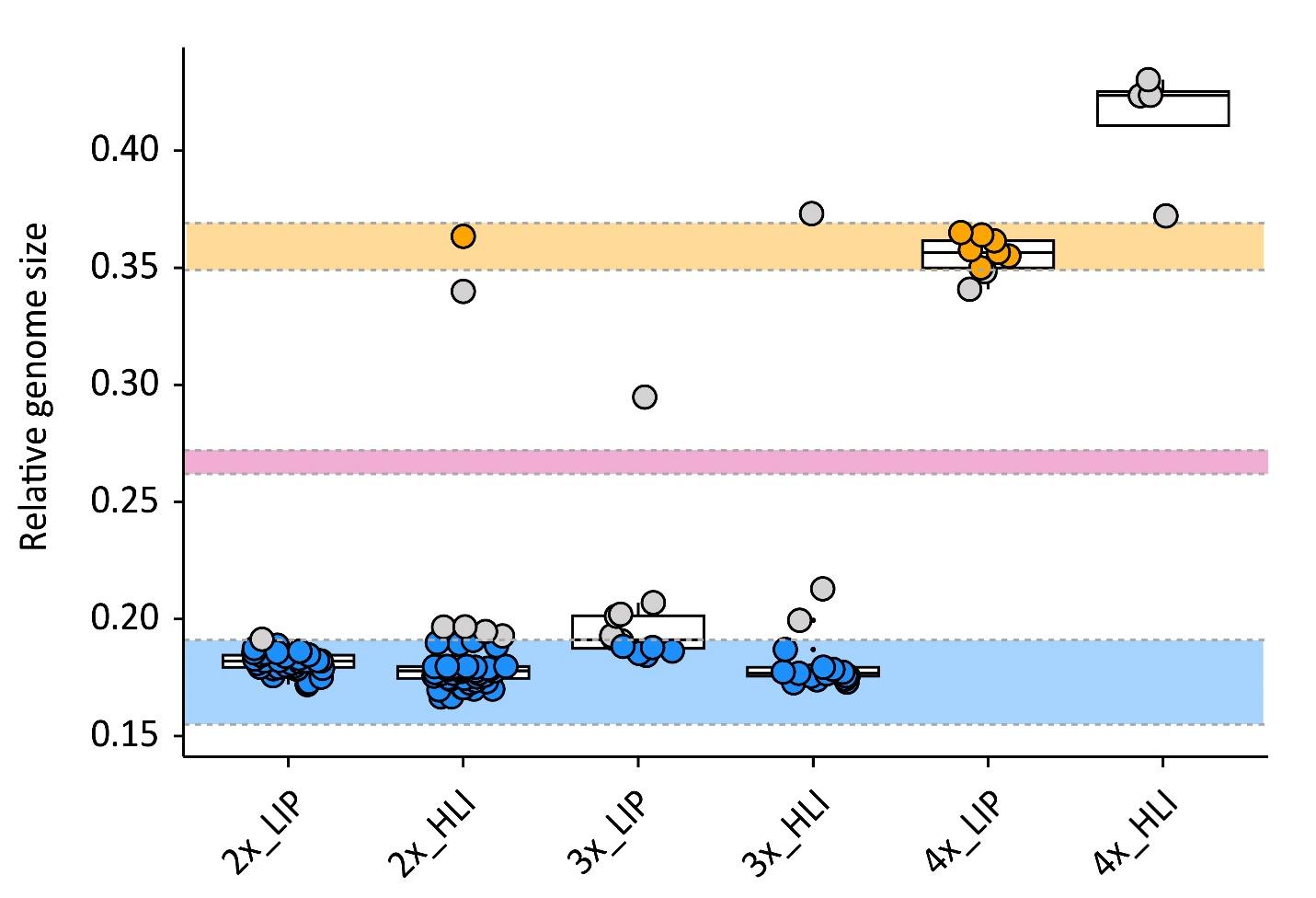

**Fig S3.** Relative genome size of seedlings germinated from seeds collected from HLI and LIP mixed ploidy populations. Out of 122 seedlings germinated, 80 were collected from diploid, 29 from triploid, and 13 from tetraploid mothers. The colour signifies the individual's ploidy: blue for 2x, pink for 3x, orange for 4x, grey symbol for aneuploids. Dashed lines represent the borders of the expected genome size ranges for 2x, 3x, and 4x individuals, respectively, defined based on a reference set of euploid individuals from the controlled homoploid crosses (see Methods for details).

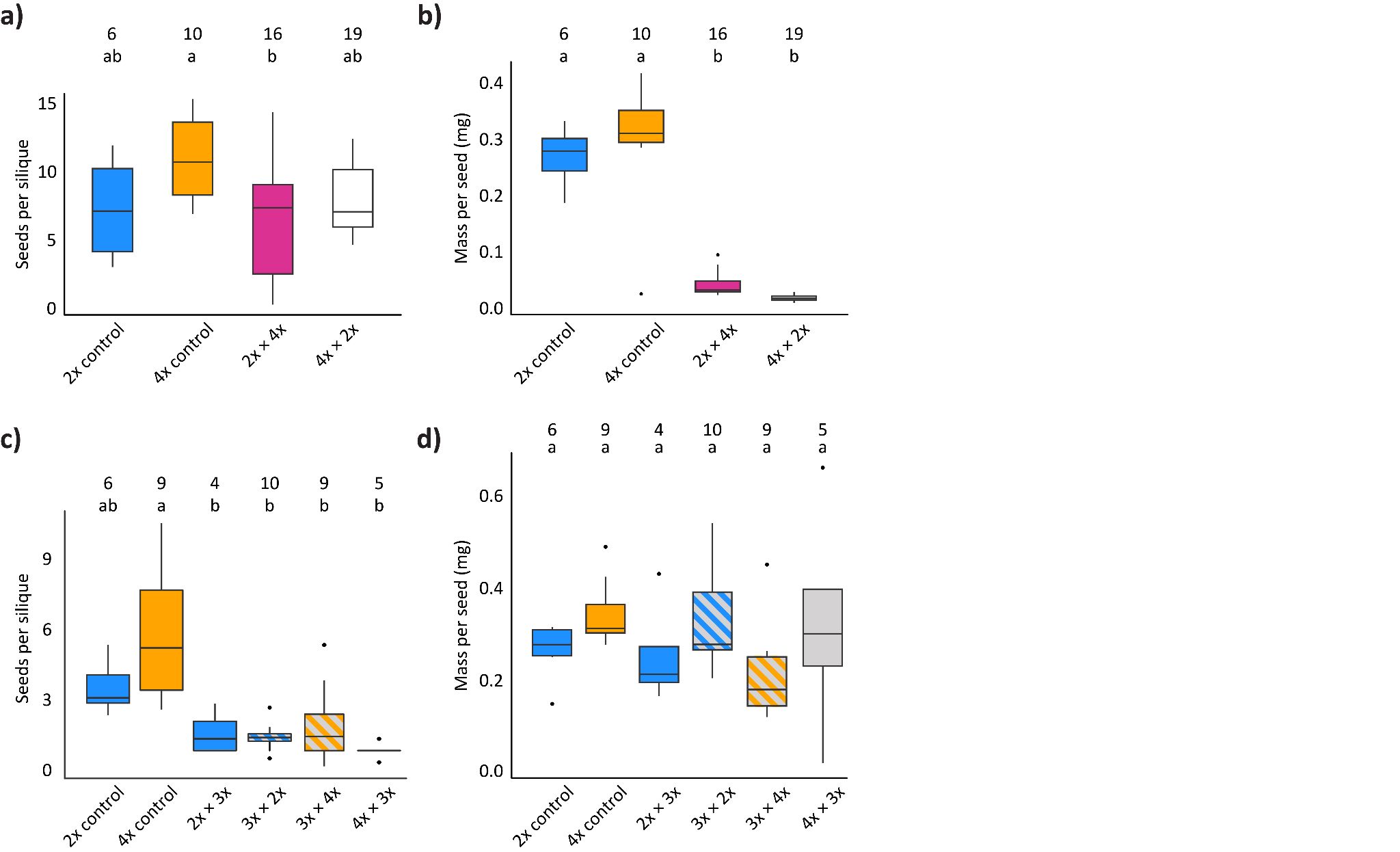

**Fig S4**. Different seed set and seed mass in the progeny of Experiment 1 (upper row a, b) and Experiment 2 (bottom row, c, d) gathered following hand pollinations of *C. amara*. (a, c) The average number of seeds per silique in successful crosses, (b, d) Average mass per seed (mg); the colours designate the obtained ploidy of the offspring: blue for 2x, purple for 3x, orange for 4x, grey for aneuploids, striped for a mixture of different cytotypes. The numbers of pollinated parental plants are shown above each bar. Letters indicate significant differences at *p* < 0.001.

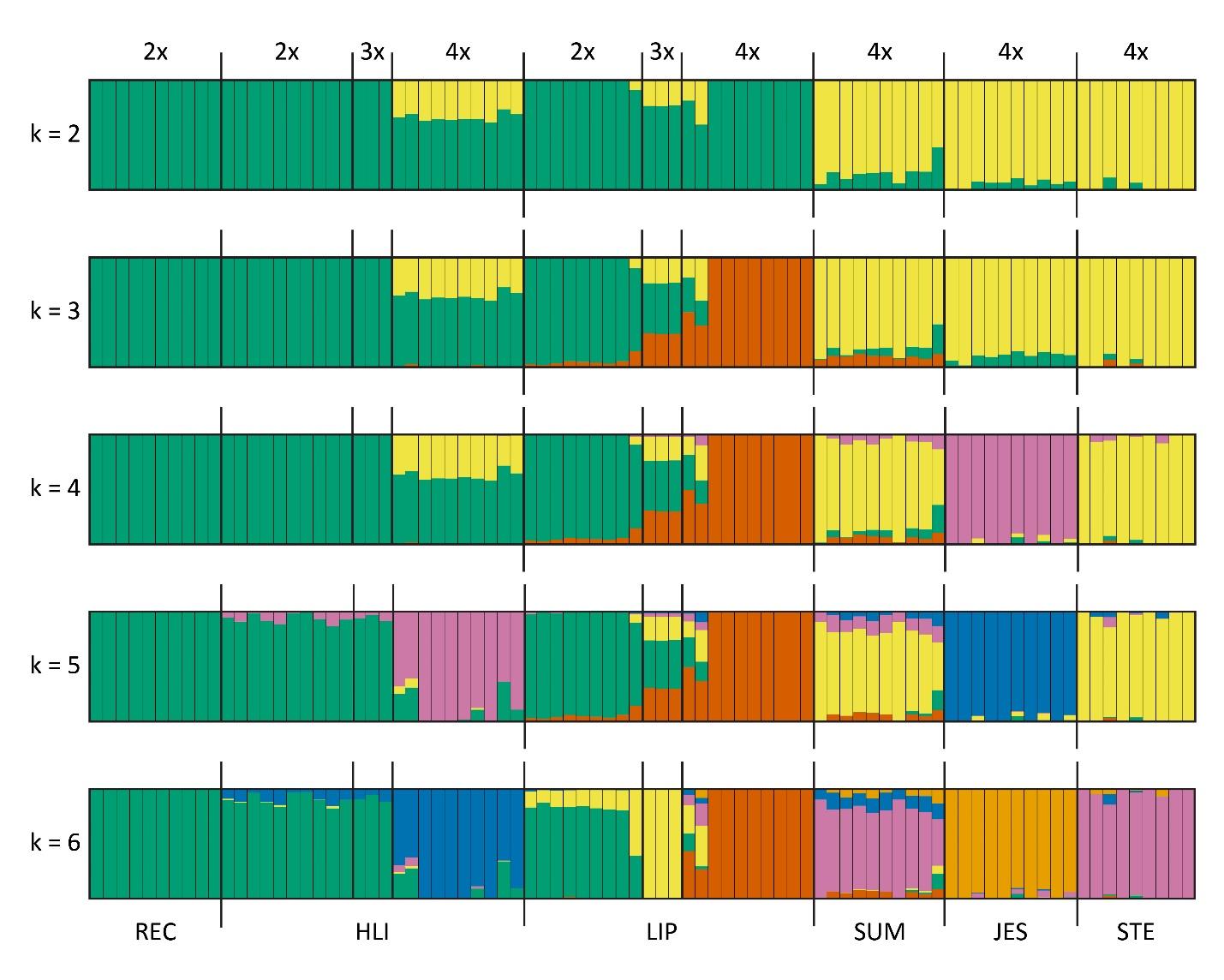

**Fig S5**. Population structure of the six populations of *C. amara* in Central Europe at K = 2 to K = 6 retrieved by STRUCTURE. The probability of assignment of each individual to a different cluster(s) is depicted by a different colour(s) within a vertical column; the populations are separated by a thicker black line with the name of the population.

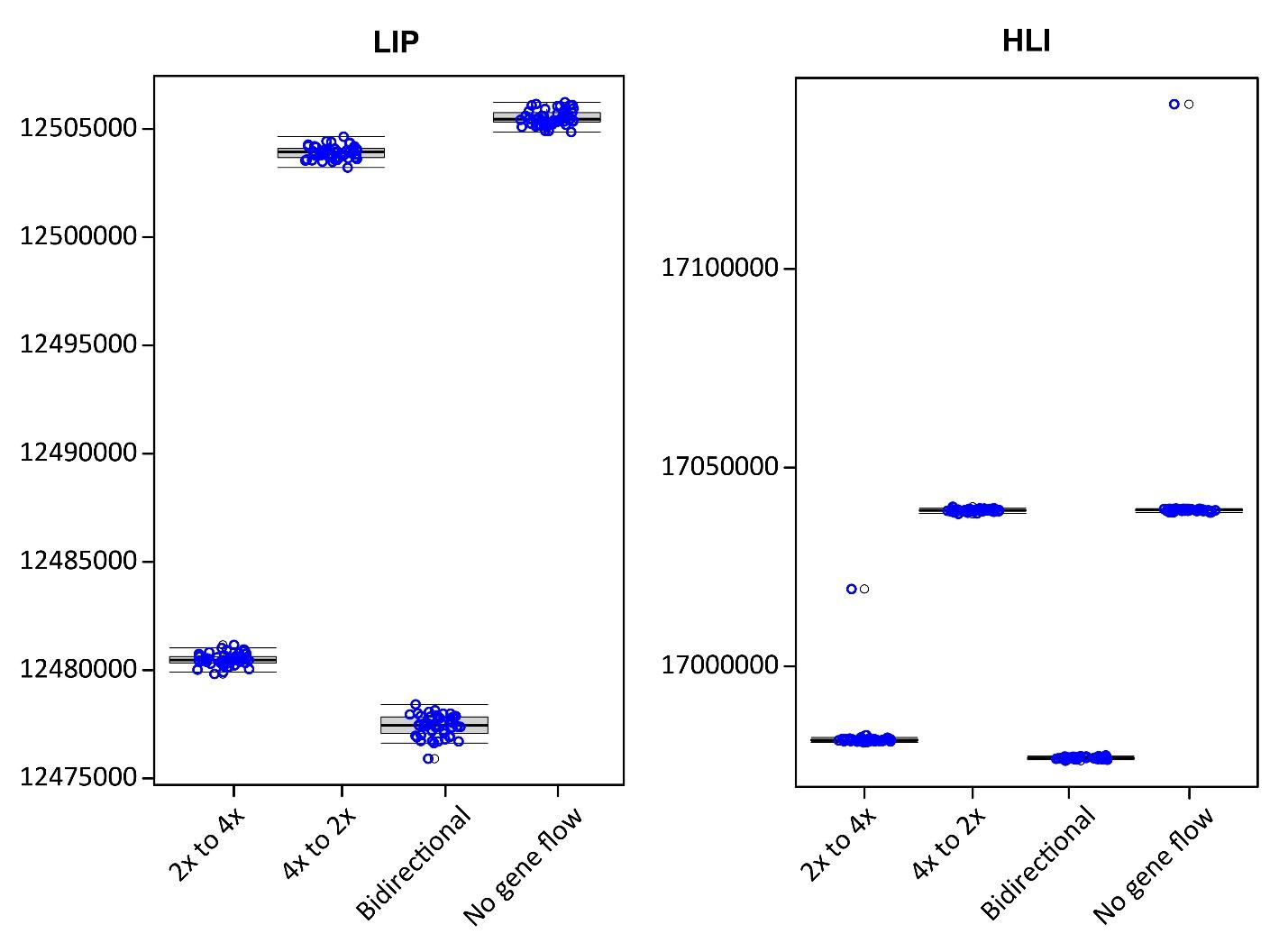

**Fig S6**. Comparison of Akaike information criteria (AIC) across four scenarios approximating the gene flow between diploids and tetraploids in mixed ploidy populations. Each scenario was simulated by 50 independent fastsimcoal runs, and the corresponding distribution of the AIC values over these 50 runs (blue circles) was summarised by the boxplots.

**Table S1** Sampling details of populations of *C amara* and average relative genome size of individuals from these populations.

| ***Population*** | ***Coordinates*** | ***Habitat*** | ***Ploidy*** | ***Number of individuals cytotyped*** | ***Mean RGS*** | ***Standard deviation ±*** | ***Number of individuals sequenced*** |
| --- | --- | --- | --- | --- | --- | --- | --- |
| **LIP** | 49.97559N, 13.24187E | Stream bank | 2x | 140 | 0,1779 | 0.1738/0.1819 | 9 |
|  |  |  | 3x | 13 | 0,2668 | 0.2652/0.2684 | 3 |
|  |  |  | 4x | 44 | 0,3577 | 0.3547/0.3608 | 10 |
| **HLI** | 49.74213N, 14.7213E | Stream bank | 2x | 79 | 0,1763 | 0.1703/0.1763 | 10 |
|  |  |  | 3x | 6 | 0,2671 | 0.2657/0.2686 | 3 |
|  |  |  | 4x | 34 | 0,3597 | 0.3569/0.3625 | 10 |
| **JES** | 50.14109N, 16.78909E | Stream bank | 4x | 40 | 0,3654 | 0.3594/0.3712 | 10 |
| **SUM** | 48.97596N, 13.77065E | Stream bank | 4x | 30 | 0,3652 | 0.3572/0.3732 | 10 |
| **STE** | 47.81211N, 14.13067E | Stream bank | 4x | 30 | 0,3693 | 0.3628/0.3759 | 9 |
| **REC** | 50.49832N, 14.90310E | Stream bank | 2x | 30 | 0,1769 | 0.1728/0.1809 | 10 |

**Table S2** Summary of crossing experiments (available in the standalone Supplementary_Tables.xlsx file).

**Table S3** Details on the genome sequenced individuals (available in the standalone Supplementary_Tables.xlsx file).

**Table S4** Relative genome sizes of putative aneuploid individuals from field samples and experimental progeny with chromosome counts.

| ***Individual*** | ***Origin*** | ***Parental ploidies*** | ***Ploidy*** | ***RGS*** |
| --- | --- | --- | --- | --- |
| **LIP_168** | field sample | ? | putative 2x-3x aneuploid | 0,197 |
| **LIP_86** | field sample | ? | putative 2x-3x aneuploid | 0,257 |
| **HLI_6** | field sample | ? | putative 3x-4x aneuploid | 0,341 |
| **F_8** | cross | 3x × 4x | hypotetraploid (2n = 31) | 0,348 |
| **M_8** | cross | 3x × 2x | hypotetraploid (2n = 31) | 0,344 |
| **L_1** | cross | 3x × 4x | hypopentaploid (2n = 38) | 0,418 |
| **D_7** | cross | 2x × 2x | diploid (2n = 16) | 0,181 |
| **B_6** | cross | 4x × 4x | tetraploid (2n = 32) | 0,357 |

**Table S5** Results of ABBA-BABA tests for introgression detection between P2 and P3.

| Tested topology | | | |  |  |  |  |
| --- | --- | --- | --- | --- | --- | --- | --- |
| **P1** | **P2** | **P3** | **O** | **D-statistic** | **jackknife** | **Z score** | **CI** |
| STE (AT 4x) | HLI (4x) | HLI (2x) | CA022 (BAL) | 0,3062 | 0,3064 | 23,885 | 0,2812 - 0,3315 |
| STE (AT 4x) | LIP (4x) | LIP (2x) | CA022 (BAL) | 0,2846 | 0,2847 | 21,094 | 0,2582 - 0,3111 |
| STE (AT 4x) | JES (4x) | LIP (2x) | CA022 (BAL) | 0,0447 | 0,0447 | 2,784 | 0,0142 - 0,0752 |
| STE (AT 4x) | SUM (CZ 4x) | LIP (2x) | CA022 (BAL) | 0,0807 | 0,0808 | 8,454 | 0,062 - 0,0995 |
| STE (AT 4x) | JES (4x) | HLI (2x) | CA022 (BAL) | 0,0541 | 0,0541 | 3,343 | 0,0224 - 0,0859 |
| STE (AT 4x) | SUM (CZ 4x) | HLI (2x) | CA022 (BAL) | 0,0801 | 0,0801 | 8,104 | 0,0607 - 0,0995 |
| STE (AT 4x) | REC (CZ 2x) | LIP (2x) | CA022 (BAL) | 0,4067 | 0,407 | 26,903 | 0,3774 - 0,4367 |
| STE (AT 4x) | REC (CZ 2x) | HLI (2x) | CA022 (BAL) | 0,4208 | 0,4211 | 25,427 | 0,3887 - 0,4536 |
| STE (AT 4x) | SUM (CZ 4x) | REC (CZ 2x) | CA022 (BAL) | 0,0775 | 0,0776 | 7,556 | 0,0574 - 0,0977 |
| STE (AT 4x) | JES (CZ 4x) | REC (CZ 2x) | CA022 (BAL) | 0,0518 | 0,0518 | 3,277 | 0,0203 - 0,0833 |
| STE (AT 4x) | SUM (CZ 4x) | JES (CZ 4x) | CA022 (BAL) | 0,2583 | 0,2584 | 29,837 | 0,2372 - 0,2972 |
